# Hydrophobic Ion Pairing for Simple, Non-Toxic Transfection

**DOI:** 10.1101/2025.04.25.650611

**Authors:** Mikaela A. Gray, Michelle Seeler, Catalina Montoya, Jaeyoung Park, Linh D. Mai, Kathryn A. Whitehead, Julie A. Champion

## Abstract

Although biomacromolecules require intracellular delivery for therapeutic effect, existing transfection agents are often characterized by high cost, low efficiency, and/or cytotoxicity. Here, we describe a new transfection approach based on hydrophobic ion pairing (HIP), which involves the simple mixing of a hydrophobic counterion with charged biomacromolecules. Among tested cargoes (proteins, siRNA, and pDNA), the HIP siRNA system performed especially well, achieving silencing in fibroblasts (80%), T cells (90%), and neurons (70%). HIP siRNA was also highly potent in mice, with tropism dependent on the route of administration. Most notably, intraperitoneal administration enabled ∼40% LAMP-1 knockdown in the pancreas, and intravenous delivery resulted in a remarkable 80% silencing in the heart. Heart delivery was also highly selectively, with no significant knockdown in the liver. Together, these data demonstrate a new, inexpensive approach to biomacromolecular delivery with the potential to target difficult-to-transfect organs, thus expanding the therapeutic potential of nucleic acids.

---

Transfection of cells with deoxyribonucleic acid (DNA), ribonucleic acid (RNA), or proteins is a critical, but often low efficiency step in research and therapeutic development. Typical methods of transfection are biological (ex. viruses), chemical (ex. nanoparticles), and physical (ex. electroporation)^1^. Of these, chemical transfection is attractive due to its efficiency across cell types, ease of use, biosafety, and cost compared to other methods. However, chemical methods face challenges of endosomal entrapment and degradation^2^.

Most chemical transfection approaches involve the complexation of a cationic agent with negatively charged cargo. These cationic molecules, which are often lipids, mitigate electrostatic repulsion by the negatively charged cell membrane and, once in the endosome, mediate escape into the cytoplasm by disrupting the negatively charged endosomal membrane^1^. One commonly used cationic lipid transfection agent is Lipofectamine™; however, its strong positive charge causes undesired toxicity that prevents its use *in vivo*^3^.

A variety of alternative nanocarriers have been developed to overcome this limitation for biomacromolecule delivery, including those created from lipids and/or polymers^4–6^. For example, lipid nanoparticles (LNPs), which are the most clinically advanced non-viral nucleic acid transfection reagent, are FDA-approved in several siRNA and mRNA products, enabling new classes of therapeutics and vaccines^7,8^. However, LNP formulations are chemically complex, typically comprising four lipids that condense RNA into nanoparticles, which complicates manufacturing. Additionally, because the particles are held together non-covalently, cold chain storage and distribution is often required to overcome stability challenges.

Alternatively, polymer particles have been widely studied as delivery carriers, with positively charged, medium length (∼10-100 kDa) polymers condensing and complexing with the nucleic acid cargo. An advance in these carriers uses hydrophobic ion pairing (HIP) to improve encapsulation efficiency by reversibly complexing the biomacromolecule with an oppositely charged hydrophobic ion, thereby increasing its association with the hydrophobic polymer^9^. The ratio of cargo to counterion determines the rate of biomacromolecule release, with higher proportions of counterion resulting in slower release. Although these polymeric systems have been shown to improve delivery compared to unpaired cargo^9–14^, they and other polymer carriers also suffer from numerous challenges, including polymer polydispersity, stability issues, and toxicity associated with higher molecular weight formulations^15^.

Here, we describe the creation of a much simpler transfection solution: HIP complexes formed through the straightforward mixing of biomacromolecule cargos with low molecular hydrophobic ions. Specifically, we paired siRNA, plasmid DNA and protein cargos with these ions and demonstrated functional cytosolic delivery to various cell types without cytotoxicity. Further, siRNA-HIP complexes effectively silenced gene expression in mice and exhibited unique tissue tropism depending on the route of administration, including to cardiac tissue.

Together, these data illustrate that HIP is a potent and non-cytotoxic delivery strategy that is generalizable to a wide variety of drug cargos, including proteins, pDNA, and siRNA.

## Results

### HIP enables non-toxic intracellular siRNA, pDNA, and protein delivery

The selected counterions in this work have one charge group and no hydrophilic head group (Figure 1 & Figure S1), making them unique from systems with multiple components such as self-emulsifying drug delivery systems, polyelectrolyte complexes, and lipid nanoparticles^10,16,17^. Only ions that are FDA approved or generally regarded as safe (GRAS) were used to maximize translational potential. We evaluated a cationic counterion, benethamine (BA), to deliver negatively charged cargos into the cytosol. BA is a lipophilic amine, used as a salt in the FDA approved formulation of long-acting penicillin G, benzylpenicillin^18^. BA has also been used to improve encapsulation efficiency of retinoic acid in nanoparticles, leading to greater anti-cancer activity than two other cationic counterions, and is non-toxic to cancer cell lines without retinoic acid^19^. Cationic lipids often have significant toxicity because the hydrophilic headgroup activates pro-apoptotic pathways and pro-inflammatory cascades in a structure dependent manner^20^. BA, however, lacks a headgroup entirely.

**Figure 1.**
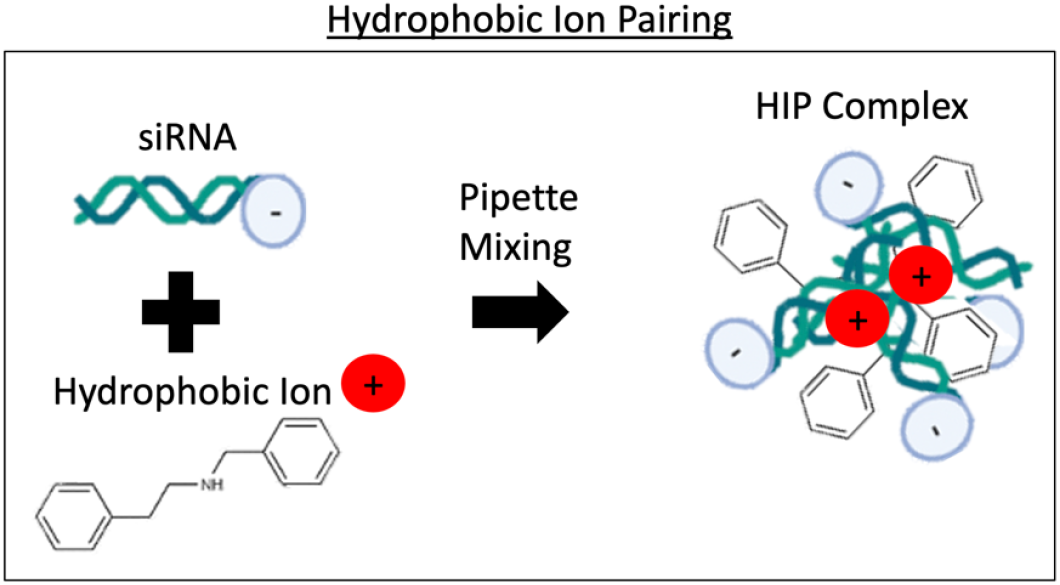
Schematic of hydrophobic ion pairing (HIP) between the positively charged hydrophobic ion benethamine (BA) and negatively charged small interfering RNA (siRNA). Figure created with Biorender.com.

To make HIP complexes of siRNA and BA, counterion was suspended in phosphate buffered saline (PBS), then mixed 2-3 times using a pipette with an equal volume of siRNA in PBS (Table S1). Distribution coefficient (LogD) was utilized to measure change in siRNA hydrophobicity upon HIP complexation, with a more positive value indicating increased hydrophobicity and distribution in butanol compared to water^21^. Multiple charge ratios were tested whereby the greatest LogD occurred at a charge ratio of 1 siRNA : 0.03 BA (Figure 2a). This charge ratio is significantly lower than that commonly used for nucleic acids and cationic lipids using LNPs, which ranges from 1 : 1 to 1 : 3^22,23^. To assess the potential of HIP complexes for functional delivery without a nanoparticle carrier, HIP complexes were formulated with anti-eGFP siRNA and BA and incubated at RT for 3 days. Then, they were applied to eGFP-expressing NIH-3T3 cells at an siRNA dose of 75 nM for 48 hours in the presence of serum containing media. Lipofectamine™ RNAiMax, a commonly used *in vitro* transfection agent, was used as a control. HIP complexing of siRNA with BA resulted in greater than 70% eGFP fluorescence knockdown (Figure 2b), similar to the Lipofectamine™ control, indicating successful cytosolic delivery. Critically, there was no cellular toxicity from the siRNA-BA complexes while Lipofectamine™ induced greater than 50% toxicity (Figure 2c), as has been reported in other cell lines^24^. Additionally, HIP complexes can be used with complete media unlike Lipofectamine™, which requires serum free media.

**Figure 2.**
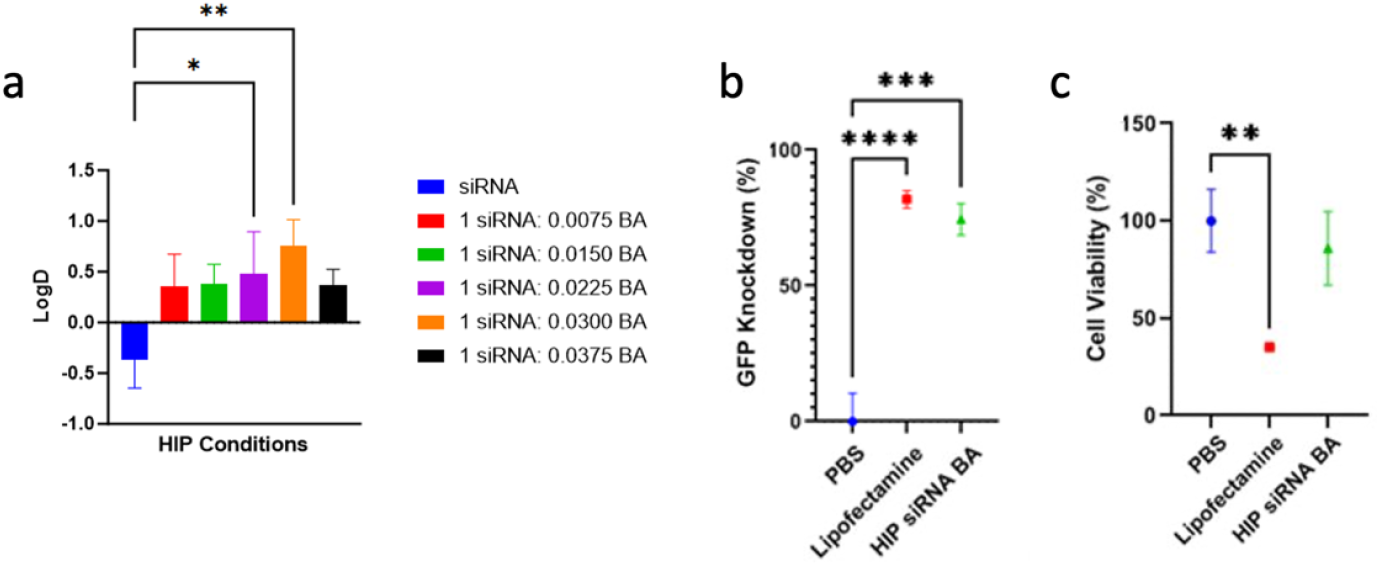
siRNA HIP complex characterization and eGFP knockdown. a) siRNA BA HIP complex hydrophobicity measured by LogD coefficient 3 days after mixing at different charge ratios. LogD was calculated based on relative siRNA concentration in hydrophobic (butanol) versus hydrophilic (water) phases. b) Quantification of eGFP knockdown in NIH3T3/eGFP cells after 48 hours incubation with 75 nM siRNA formulated in 1 siRNA: 0.03 BA charge ratio HIP complexes and Lipofectamine™ RNAiMAX, or PBS only control. Cell media contained 10% FBS for all groups except Lipofectamine™. Live cell populations were selected after adding trypan blue to trypsinized cells using flow cytometry. c) Quantification of cellular viability as measured by metabolism using 3-(4,5-dimethylthiazol-2-yl)-2,5-diphenyltetrazolium bromide(MTT) assay. Influence on % cell viability after 48 hours was normalized to PBS treated cell viability. * p < 0.05, ** p < 0.01, *** p <0.001, and **** p < 0.0001.

Currently, LNPs are considered as a clinical gold standard carrier for RNA delivery. To compare HIP and lipid nanoparticles, we formulated LNPs with the ionizable lipid 306O_10_^25^ and delivered anti-eGFP siRNA to NIH-3T3/eGFP cells at various siRNA concentrations using 1 siRNA: 0.03 BA HIP complexes immediately after mixing. LNPs and HIP complexes performed comparably from 74.8-18.7 nM siRNA doses with 70-80% GFP knockdown after 48 hours indicating the robustness of HIP at delivering siRNA (Figure 3a-b). Additionally, HIP complexes showed a slight reduction in viability at the highest dose while lipid nanoparticles at all concentrations had a slight reduction in viability (Figure 3c).

**Figure 3.**
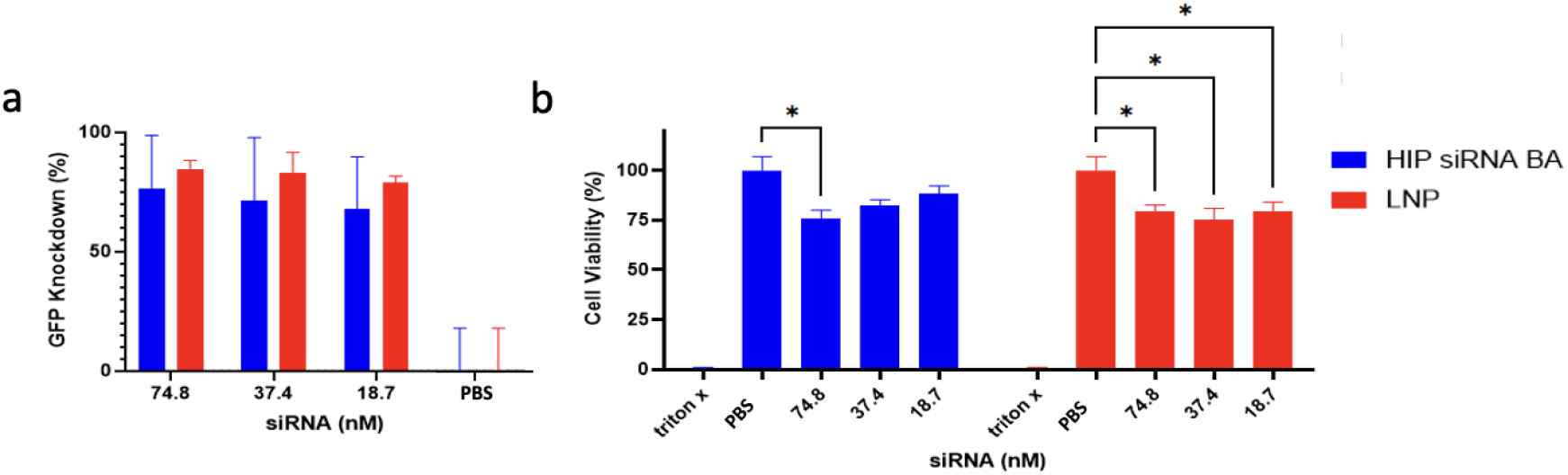
HIP siRNA BA and LNP delivery and viability comparison. Quantification of eGFP knockdown (a) in NIH3T3/eGFP cells after 48 hours incubation with 1 siRNA: 0.03 BA HIP complexes (blue) or LNP (red) at varying doses of siRNA. Cell media contained 10% FBS for all groups. All formulations containing siRNA are significantly different from the untreated control (p < 0.0001) b) Quantification of cellular viability as measured by metabolism using 3-(4,5-dimethylthiazol-2-yl)-2,5-diphenyltetrazolium bromide(MTT) assay. Cell viability was normalized to untreated control. * p < 0.05.

After delivering the double stranded 21-mer siRNA HIP complexes, we examined delivery of a much larger double stranded cargo, 5.4 kilobase pair (kbp) plasmid DNA (pDNA), by BA HIP. To make HIP complexes, BA was diluted in PBS, then mixed 2-3 times with a pipette with an equal volume of pDNA to achieve various charge ratios (Table S1). To identify the optimal amount of counterion for pDNA delivery, 2 µg per 96-well of 15,000 cells of a luciferase encoding plasmid complexed with BA at a range of charge ratios (1 pDNA:3.2 × 10^−7^ to 3.2 × 10^−6^ BA) was incubated with HeLa cells immediately after mixing or three days after mixing and luciferase expression was evaluated after 72 hours (Figure S2). The ratios are much lower than for siRNA because the size and charge of pDNA is much larger than siRNA but BA is not soluble in PBS above 15 mg/mL. Only one ratio, 1 pDNA : 1.6 ×10^−6^ BA resulted in functional delivery as BA. Higher charge ratios were not effective, likely because the BA was not soluble, as precipitation was observed. While transfection efficiency was much lower than for Lipofectamine™ 2000, HIP can be dosed in the presence of serum, and does not require refrigeration.

Given that HIP has been performed with some peptides and model proteins^9^, we also investigated protein delivery by HIP complexes using cytochrome c (CytC) as a model therapeutic since it a cytosolically active protein that induces apoptosis when in the cytosol^26^. Two GRAS anionic counterions, oleic acid (OA) and sodium docusate (SD), were tested with cationic CytC as well as cationic sfGFP, as a non-toxic control cargo. OA is a fatty acid and common ingredient in cholesterol supplements that has been used to improve the encapsulation efficiency of insulin, polymyxin B, and others in nanoparticle delivery systems^9,27–29^.

Additionally, OA acts as a permeation enhancer for transdermal nanoparticle delivery by reducing skin barrier function and improving permeation of drugs.^30^ Drugs that use OA to improve skin permeation include celecoxib, adapalene, and tranilast for the treatment of pain, acne, keloids, and other conditions^30–33^. SD is the most commonly used ingredient in laxatives. It is the second most common anionic counterion used behind sodium dodecyl sulfate for improved nanoparticle encapsulation, but has less safety concerns and is a highly effective counterion for increasing encapsulation efficiency^14,34^.

Protein HIP complexes were made at a range of charge ratios. Pairing with counterions did not largely affect the structure of CytC or sfGFP(+10), as indicated by retention in absorbance and fluorescence, respectively, though OA complexation at a high charge ratio did reduce sfGFP(+10) fluorescence by 24.2 ± 0.6 % (Fig S3a-c). Complexes exhibited a particulate-like nature (Fig S3d-f). Delivery was assessed in HeLa cells in serum containing media for 48 hours of incubation. Cytotoxicity was used as the indicator of successful cytosolic delivery of CytC. Significantly decreased cell viability occurred at charge ratios above 1 CytC: 10.66 counterion for both OA and SD (Fig S3g). Beyond a charge ratio of 1 CytC: 16 counterion there was no additional delivery, or cytotoxicity, benefit. Delivery of non-cytotoxic protein control, cationic sfGFP(+10), by OA and SD HIP complexes showed no reduction in viability (Fig S3h), demonstrating that HIP-protein complexes are non-toxic and the delivered CytC was the cause of decreased viability (Fig S3g). However, the dose of CytC required to kill cells was relatively high, 20 µM, compared to 2 µM needed for HIP loaded protein vesicles^35^. Further, the charge ratios required for protein delivery were orders of magnitude higher than for nucleic acids, indicating that protein delivery by HIP complexes is less efficient than nucleic acid delivery.

This may be due to the zwitterionic nature of proteins and their hydrophobic core, into which hydrophobic ions may partition.

### HIP siRNA BA silences hard to transfect cells

Since siRNA delivery using HIP was efficient, we selected this cargo to evaluate delivery to traditionally hard-to-transfect cell types. To measure gene silencing, anti-lysosomal-associated membrane protein 1 (LAMP1) siRNA was selected as LAMP1 is expressed widely across cell types in the plasma membrane, in addition to lysosomal membranes, making its expression easy to assess with antibodies^36^. T cells are difficult to transfect due to their ability to restrict viral replication^37^. However, they are an important DNA/mRNA transfection target for FDA approved chimeric antigen receptor (CAR) T cell therapies against various leukemias^38–41^ and there is interest in delivery of miRNA to T cells^42^. Typically CAR T cells are transfected by electroporation with about 60-70% transfection efficiency^43,44^. A dose of 75 nM siRNA in BA HIP complexes induced 93.0 ± 2.2% knockdown in Jurkat human T cells, outperforming Lipofectamine™ RNAiMAX (Figure 4 & Figure S4). Neurons are difficult to transfect because they are mostly post-mitotic cells that do not divide and are highly sensitive to changes in culture conditions used for transfection^45^. HIP surpassed Lipofectamine™ RNAiMAX in HCN2 human neuronal cells with 72.5 ± 23.5% knockdown. Stem cells, commonly derived from bone marrow for regenerative medicine applications, are also hard to transfect^46,47^. HIP silenced 78.5 ± 12.0% of LAMP1 signal in primary murine bone marrow cells, comparable to Lipofectamine™ RNAiMAX.

**Figure 4.**
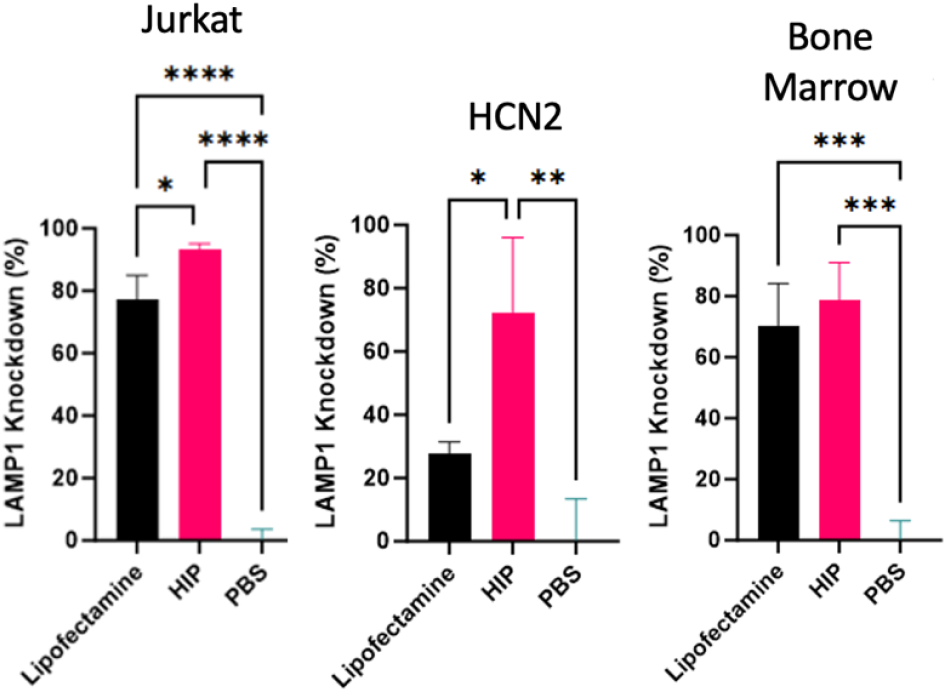
Gene silencing in Jurkat T cells (human), HCN2 neurons (human), and primary bone marrow cells (mouse) using either human or mouse anti-LAMP1 siRNA. 75 nM siRNA was given in the presence of FBS for all groups except for Lipofectamine™ RNAiMAX and treated for 48 hours. Gene silencing was measured by labelling cells with either human or mouse anti-LAMP1 antibody and measuring mean fluorescence using flow cytometry. LAMP1 knockdown is relative to unsilenced PBS control fluorescent signal. * p < 0.05, ** p < 0.01, *** p < 0.001, and **** p < 0.0001.

### *In vivo* administration of HIP siRNA BA using intraperitoneal, intravenous, and intranasal routes shows organ tropism

Given the *in vitro* capability of HIP delivery of siRNA, we evaluated it for *in vivo* delivery in mice using the same 1 siRNA: 0.03 BA charge ratio. Anti-LAMP1 siRNA was selected as LAMP1 is ubiquitously expressed across cell types and organs and silencing LAMP1 has no effect on mouse health^36^. Administration route influences nucleic acid loaded LNP organ distribution, expression kinetics, and therapeutic outcomes^7,48,49^. Thus, we selected three systemic routes of administration: intraperitoneal (IP), intravenous (IV), and intranasal (IN) and compared gene silencing in the liver along with specific organs of interest for each route as most nanoparticles accumulate in the liver^6^. We selected organs of interest and siRNA dose for each route out of all collected organs (all routes: heart, liver, spleen, pancreas, kidneys; IP: small intestine; IV: bone marrow; IN: brain) from a preliminary experiment that evaluated 1.5, 3, and 5 mg/kg siRNA dosages (Figures S5-7). From this experiment, doses selected were 1.5 mg/kg for IP and IN and 5 mg/kg for IV, and cells from organs were collected 48 hours after administration. Negative controls consisted of saline and scrambled siRNA complexed by HIP with BA at 1:0.03 charge ratio.

IP administration induced 49.0 ± 16.3 % LAMP1 knockdown in the pancreas, with no liver LAMP1 knockdown (Figure 5a, Figure S8a). As a point of comparison, Melamed and coworkers created an ionizable LNP loaded with mRNA, injected IP, that had greater than 60% delivery specificity for the pancreas with some delivery to the spleen and liver^50^. In the case of HIP, IV administration via the jugular vein showed a remarkable 84.9 ± 5.9 % LAMP1 knockdown in the heart with no functional delivery detectable in any other organs (Figure 5b, Figure S8b). Both the high potency in the heart and the negligible transfection of the liver is highly unusual for RNA delivery systems, and to our knowledge, such effective and selective heart siRNA delivery has not been previously reported^51–53^. These unique effects may derive from the absence of a typical nanocarrier, as is suggested by a study in which HIP-like complexes of the ionizable amine fulvestrant drug and siRNA loaded into LNPs resulted in only liver knockdown when administered IV^54^. IN administration resulted in 59.9 ± 14.1 % knockdown in the liver and 59.4 ± 10.3 % knockdown in the heart (Figure 5c, Figure S8c). IN delivery to the heart is an emerging approach for rapid drug absorption through arteries in the respiratory area that drain into the internal jugular veins and into the heart through the superior vena cava^55^. We hypothesize that both IN and IV routes resulted in cardiac delivery since both either start in or quickly reach the jugular veins. There were no significant changes in mouse weight over 48 hours, indicating no gross impacts on animal health (Figure S9).

**Figure 5.**
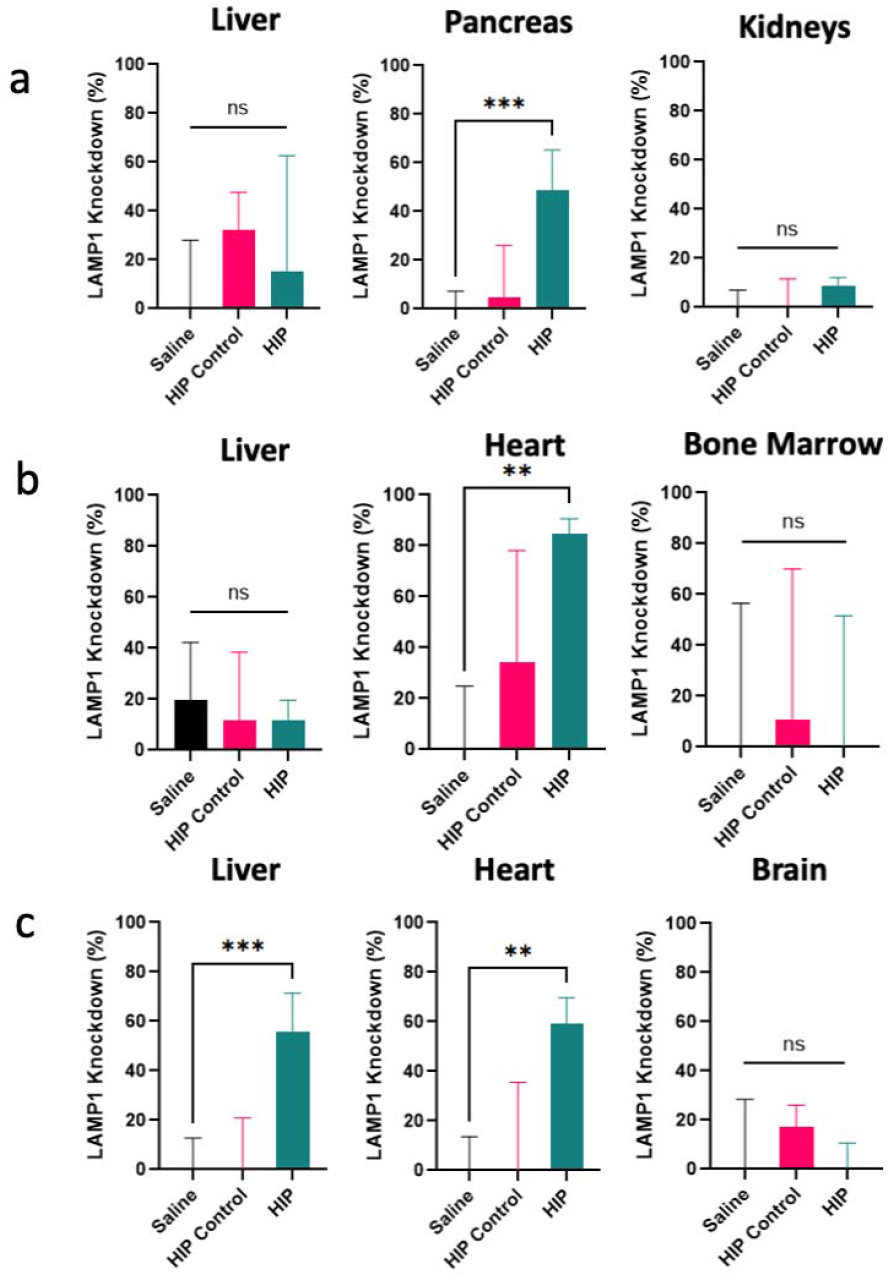
*In vivo* biodistributions using a) IP, b) IV, and c) IN routes in (n=5) BALB/c mice each using an anti-LAMP1 siRNA dose of 1.5 mg/kg, 5 mg/kg, and 1.5 mg/kg. Organs were homogenized, red blood cells were removed, and an anti-LAMP1 antibody was used to assess LAMP1 gene silencing using flow cytometry. LAMP1 knockdown single cell population values 48 hours after treatment. One-way ANOVA and the F test was used to compare groups for statistical significance. Outliers were removed (from saline group only as experiments were performed on different days) using Grubbs’ statistical test. * p < 0.05, ** p < 0.01, and *** p < 0.001.

## Discussion

Simple, efficient, and non-toxic delivery is essential for widespread use of nucleic acids in research and therapeutics. Current methods are complex and require formulation optimization with multiple chemical components^3,56^ or specialized equipment like electroporators^43,44^. These results demonstrate that HIP overcomes these challenges and has the potential to broaden the use of nucleic acids to non-experts or those with fewer resources. Using pipette mixing of counterion with biomacromolecule, we report cytosolic delivery of siRNA, pDNA, and protein. HIP delivery was demonstrated in six different cell types, including hard-to-transfect cells. siRNA HIP delivery is as effective as LNPs *in vitro* at the concentrations tested, and as effective or more effective than Lipofectamine™ RNAiMAX, the commercial gold standard, without negative impact on cell viability. At the current price of $335 per 100 g, BA is 14 million times cheaper per transfection reaction than Lipofectamine™ RNAiMAX, with a current price of $160 per 100 µL, and does not require refrigeration. HIP pDNA delivery was 6,756 times cheaper than ExpiFectamine™ when normalized to expression output, with a current price of $916 per L of culture while HIP pDNA delivery of a 5.4 kbp plasmid currently costs $0.08 per L of culture.

In hard-to-transfect T-cells, HIP outperformed Lipofectamine™ RNAiMAX with 90% LAMP1 silencing using 75 nM siRNA. CAR T cell therapy faces the challenges of resistance and toxicity due to incidental lentivirus integration^57^. To mitigate these challenges and improve clinical outcomes of CAR T cell therapy, researchers are evaluating the use of short hairpin RNA in three on-going clinical trials^58–60^. We postulate that HIP is a feasible, easy, and cheap way for implementation of small RNAs in CAR T cell therapeutics. Additionally, we hypothesize that HIP could work well for other oligonucleotides such as antisense oligonucleotides or doggybone™ DNA.

A major limitation of nanoparticle therapeutics is liver biased delivery. Since particles typically accumulate in the liver, therapeutic applications have focused on gene targets in the liver. By varying the route of injection, HIP formulated siRNA can be utilized to give liver or non-liver delivery. HIP complexes also deliver to hard-to-deliver to organs such as the pancreas and heart. Cardiac delivery of HIP formulations using a less invasive injection route (IV or IN) could expand the field of nucleic acid development into more disease applications. Most LNPs targeting cardiovascular diseases are injected intramyocardially to localize cargo in the cardiac tissue^61–64^, a route that requires open-chest surgery. In the future, other systemic and local routes of siRNA delivery by HIP complexes can be evaluated to identify other organs and diseases with translational potential. The simplicity and low cost of HIP formulations make it a valuable tool in screening libraries of silencing targets for a variety of diseases both *in vitro* and *in vivo*.

Altogether, the results presented here demonstrate the potential for nucleic acid HIP complexes to be applied widely for drug discovery, basic research, therapeutics manufacturing, and clinical use.

## Methods

### Bacterial Protein Expression & Purification

The pET28a-sfGFP-His plasmid with cationic sfGFP(+10) variant^65^ was purchased from Genscript. sfGFP variants were expressed in *Escherichia coli* strain BL21 (DE3) star. To express these proteins, 1 L of lysogeny broth containing 50 mg kanamycin was inoculated with 5 mL of overnight culture at 37°C then induced with 1 mM isopropyl ß-D-1 thiogalactopyranoside when the optical density was greater than 0.7. After 5 hours of expression, cultures were collected by centrifugation at 4000 g for 10 minutes. The pellets were resuspended in lysis buffer containing 300 mM NaCl, 50 mM NaH_2_PO_4_, and 10 mM imidazole and lysed by sonication. Next, the lysate was cleared by centrifugation at 10,000 g for 10 minutes and incubated with Ni-nitrilotriacetic acid (NTA) agarose resin (Qiagen) for at least 1 hour at 4°C. This mixture was applied to an Econo-Column (Biorad) and washed with 100 mL of lysis buffer containing 25 mM imidazole and eluted with 10 mL of lysis buffer containing 250 mM imidazole. Purity was verified using sodium dodecyl sulfate polyacrylamide gel electrophoresis (SDS-PAGE), then the proteins were buffer exchanged into phosphate buffered saline (PBS) by dialysis with three buffer exchanges at 4°C, including an overnight incubation.

### Hydrophobic Ion Paired Complex Formation and Characterization

HIP complexes were formed by simple pipet mixing. Cargo at a concentration of 1 mg/ml for proteins, 50 µM for siRNA, or 1 mg/mL for pDNA in PBS (proteins) or nuclease-free water (nucleic acids) were mixed with varying concentrations of SD, OA, or BA in PBS at equal volumes. If incubating for longer than immediate use, the complexes were left at room temperature and protected from light because many counterions are light sensitive. LogD experiments were performed by mixing 0.5 mL of water with 0.5 mL butanol then adding 100 µL cargo and HIP complexes with 0.5 mg/mL siRNA cargo. After mixing at 150 rpm overnight and phase separation, the concentration of siRNA in each phase was measured by absorbance at 260 nm (NanoDrop). LogD values were calculated by taking the log of the ratio of concentration in the butanol phase over the concentration in the water phase.

Transmission electron microscopy (TEM) grids were prepared by dropping 10 µL CytC HIP complex sample on copper grids, letting the sample adhere for 5 minutes, washing in water for 30 seconds, staining with 1% phosphotungstic acid for 10 seconds, washing again for 30 seconds, and drying overnight. A JEOL 100 CX-II TEM was utilized to visualize samples.

### Cell Culture & Delivery

HeLa cells (ATCC; taken from Henrietta Lacks, http://henriettalacksfoundation.org/) were seeded at 15,000 cells/well using Dulbecco’s modified Eagle medium (DMEM) supplemented with 10% fetal bovine serum (FBS) in 96 well plates for 24 hours prior to treatment, washed with PBS, then treated with HIP complexes (20 µM protein or 1 mg/mL pDNA) or controls at 37°C in a humidified environment containing 5% CO_2_ for 24 or 48 hours. NIH 3T3 fibroblast cells expressing eGFP (Cell BioLabs) were seeded at 35,000 cells/well in 96 well plates for 24 hours before treatment using DMEM supplemented with 10% FBS. All groups besides Lipofectamine™ were treated with DMEM and 10% FBS for a final well concentration of 75 nM siRNA. Lipofectamine™ controls followed manufacturer instructions. After incubation for 24 hours, the media was changed to give all groups 10% FBS and cultured for another 24 hours at 37°C in a humidified environment containing 5% CO_2_. HCN2 neuron cells were cultured with the same conditions as HeLa cells. Bone marrow and Jurkat cells were cultivated in Roswell Park Memorial Institute 1640 Medium (RPMI; Corning) supplemented with 10% v/v fetal bovine serum (FBS, heat inactivated; Gibco).

To assess functional delivery of anti-eGFP siRNA, NIH 3T3 eGFP cells were detached using trypsin and resuspended in PBS with trypan blue to quench any extracellular fluorescence. The eGFP fluorescence of 10,000 cells was measured on the CytoFLEX flow cytometer (BD Bioscience). Gating was done on PBS groups to isolate single cells and compare increase or decrease in eGFP signal.

To assess functional delivery of pDNA, luciferase encoding pDNA was used. HeLa cells were seeded at 15,000 cells/well in 96 well plates for 24 hours prior to treatment, then treated with HIP complexes (equal volume pDNA (2µg) and counterion at varying charge ratios) or Lipofectamine™ 2000 at 37° C in a humidified environment containing 5% CO_2_ for 72 hours.

HIP groups were treated in DMEM media with 10% FBS, while Lipofectamine™ was treated in DMEM media without FBS. After 24 hours, the Lipofectamine™ group’s media was exchanged for media containing FBS. All groups received fresh media after 48 hours. A luciferase assay was performed, after removing media and washing with PBS, by adding 100 microliters of Bright-Glo™ reagent (Promega) per well for 2 minutes then measuring luminescence using a plate reader.

To assess functional delivery in hard to transfect cell types, human and mouse anti-LAMP1 siRNA was used. Jurkat, HCN2, and bone marrow cells were seeded at 300,000 cells/well in 96-well plates 24 hours prior to treatment. HIP groups were treated in DMEM media with 10% FBS, while Lipofectamine™ RNAiMAX was treated in DMEM media without FBS. After 24 hours, the Lipofectamine™ media was exchanged for media containing FBS. After treating cells for 48 hours, bone marrow and HCN2 cells were trypsinized. Cells were blocked with Trustain FcX plus blocking solution on ice for 10 min, then stained with fluorescently labelled anti-LAMP1 (either mouse or human). HCN2 cells only were fixed with 3.7% formaldehyde for 15 minutes and permeabilized with 1% Triton X-100 for 10 minutes prior to blocking and staining to enable access to intracellular LAMP1 as extracellular LAMP1 signal was insufficient. After staining for an hour in the dark, cells were washed with 1% BSA and Jurkat and bone marrow cells were fixed with 3.7 % formaldehyde in PBS for 15 min. Cells were washed and flow cytometry analysis was conducted using a Cytoflex S flow cytometer.

### Cell Viability Assay

*In vitro* delivery of CytC and toxicity of HIP complexes were quantified using a methylthiazoltetrazolium (MTT) assay. After treatment with cargo-HIP complexes as described above, the media was removed, cells were washed with PBS, then replaced with fresh media with 10 µL of 5 mg/ml MTT solution (Biotium). The cells were incubated for 4 hours at 37°C in a humidified environment containing 5% CO_2_. Then, 200 µL of dimethyl sulfoxide (DMSO) was added to solubilize formazan crystals. Absorbance was measured to remove background and obtain viability (570 nm and 630 nm). Cell viability was calculated by dividing the number of viable cells in a treated group relative to the PBS control group.

### Lipid Nanoparticle Formulation, Delivery, & Cell Viability

LNPs were generated as previously described^25^. LNPs were formulated with the ionizable lipidoid 306O_10_, which was synthesized using Michael addition chemistry by reacting the amine 3,3′-Diamino-N-methyldipropylamine (306) with the alkyl acrylate n-Decyl Acrylate at a stoichiometric ratio of 1:4. The reaction was carried out without solvent while stirring in a 7 ml scintillation vial at 90 °C for 3 d. The lipidoid 306O_10_, DSPC (Avanti Polar Lipids), cholesterol (Sigma Aldrich), and C14-PEG2000 (Avanti Polar Lipids) were dissolved in a solution of 90% (v/v) ethanol and 10% (v/v) 10 mM sodium citrate) and combined at a molar ratio of 50:10:38.5:1.5. In a separate tube, the siRNA was dissolved in sodium citrate 10 mM. Nanoparticles were formed by rapid pipet mixing of an equal volume of the RNA and lipid solutions. The final weight ratio of lipidoid : siRNA was 5:1. Finally, nanoparticles were diluted in PBS to achieve the desired concentrations.

NIH 3T3 eGFP cells were cultured as above. Cells were seeded at 25,000 cells per well in black 96-well plates with clear, cell-culture treated bottoms (Corning). Cells were then incubated with LNPs or HIP formulation (1 siRNA:0.03 BA charge ratio) for 48 hours at siRNA doses between 25 and 200 ng. After 48 hours, eGFP fluorescence was measured at 470/515 nm using a Synergy H1 microplate reader (BioTek Instruments). Cell viability was measured as described above.

### Delivery of LAMP1 siRNA to Mice

Animal experiments were performed in accordance with the regulations and guidelines of the NIH Guide for the Care and Use of Laboratory Animals, and all protocols and procedures were reviewed and approved by Georgia Tech’s Institutional Animal Care and Use Committee (A100616). Mouse anti-LAMP1 siRNA and scrambled control siRNA were formulated by HIP with BA as described above (1 siRNA: 0.03 BA charge ratio). 6-8 week-old BALB/c female mice (Jackson Laboratory) were injected IP, IV in the jugular vein, or IN by dropwise alternating right and left nostril adding 50 µL of HIP complexes (1.5, 5, and 1.5 mg/kg siRNA, respectively) in pharmaceutical grade saline for each route. Animals were monitored for weight loss and signs of lethargy from injection until euthanasia at 48 hours.

Organs (heart, liver, pancreas, brain, kidneys, spleen, bone marrow, and small intestine) were harvested, triturated gently, and strained through 70 µm strainers. Cells were then resuspended into RMPI 1640 medium supplemented with 10% FBS and incubated with ACK red blood cell lysis buffer for 10 min. In a 96 U-bottom well plate, 1 × 10^6^ cells were seeded in each well and blocked with Trustain FcX plus blocking solution on ice for 10 min. Cells were labelled with anti-LAMP1 APC-eFluor™ 780 (ThermoFisher, 0.35 µL per well). After staining for 1 hr in the dark, cells were washed with PBS containing 1% bovine serum albumin (BSA) and fixed with 100 μL of 3.7 % formaldehyde in PBS for 15 min. Cells were washed and flow cytometry analysis was conducted using a Cytoflex S flow cytometer.

### Statistics

One-way ANOVA was used to compare 3 or more groups by using the F test for statistical significance by comparing multiple means by calculating error between all comparisons. When less than 3 groups were compared, a T-test was used for an individual two-way comparison.

## Acknowledgements

The authors acknowledge financial support from the National Science Foundation BMAT Award 2104734 and Georgia Institute of Technology Office of Technology Licensing. This work was performed in part at the Georgia Tech Institute for Electronics and Nanotechnology, a member of the National Nanotechnology Coordinated Infrastructure, which is supported by the National Science Foundation (Grant No. ECCS-2025462). In addition, this work was performed using core facilities from the Department of Animal Resources at Georgia Tech. We thank Dr. Richard Noel and veterinary staff for their assistance, support and training. We acknowledge the contributions of named and unnamed people whose health, lives, livelihoods, legacy, and privacy were extorted, often without compensation, consent, or regard to their safety, in the name of biomedical research. These men, women, and children were stripped of their humanity, and often their identity. We commit to educating ourselves and others on the history and ethical failures of biomedical research, expressing our gratitude, and encouraging others to do the same.

## References

1. Fus-Kujawa, A. et al. An Overview of Methods and Tools for Transfection of Eukaryotic Cells in vitro. Frontiers in Bioengineering and Biotechnology 9, (2021).

2. Mendes, B. B. et al. Nanodelivery of nucleic acids. Nature Reviews Methods Primers 2, 24 (2022).

3. Wang, T., Larcher, L. M., Ma, L. & Veedu, R. N. Systematic Screening of Commonly Used Commercial Transfection Reagents towards Efficient Transfection of Single-Stranded Oligonucleotides. Molecules 23, (2018).

4. Navarro, G. et al. P-glycoprotein silencing with siRNA delivered by DOPE-modified PEI overcomes doxorubicin resistance in breast cancer cells. Nanomedicine (Lond) 7, 65–78 (2012).

5. Haley, R. M. et al. Lipid Nanoparticle Delivery of Small Proteins for Potent In Vivo RAS Inhibition. ACS Appl. Mater. Interfaces (2023) doi:10.1021/acsami.3c01501.

6. Mitchell, M. J. et al. Engineering precision nanoparticles for drug delivery. Nature Reviews Drug Discovery 20, 101–124 (2021).

7. Hou, X., Zaks, T., Langer, R. & Dong, Y. Lipid nanoparticles for mRNA delivery. Nature Reviews Materials 6, 1078–1094 (2021).

8. Hu, B. et al. Therapeutic siRNA: state of the art. Signal Transduction and Targeted Therapy 5, 101 (2020).

9. Ristroph, K. D. & Prud’homme, R. K. Hydrophobic ion pairing: encapsulating small molecules, peptides, and proteins into nanocarriers. Nanoscale Adv. 1, 4207–4237 (2019).

10. Phan, T. N. Q., Shahzadi, I. & Bernkop-Schnürch, A. Hydrophobic ion-pairs and lipid-based nanocarrier systems: The perfect match for delivery of BCS class 3 drugs. Journal of Controlled Release 304, 146–155 (2019).

11. Gaudana, R., Khurana, V., Parenky, A. & Mitra, A. K. Encapsulation of Protein-Polysaccharide HIP Complex in Polymeric Nanoparticles. Journal of Drug Delivery 2011, 458128 (2011).

12. Chamieh, J. et al. Peptide release from SEDDS containing hydrophobic ion pair therapeutic peptides measured by Taylor dispersion analysis. International Journal of Pharmaceutics 559, 228–234 (2019).

13. Lobovkina, T. et al. In Vivo Sustained Release of siRNA from Solid Lipid Nanoparticles. ACS Nano 5, 9977–9983 (2011).

14. Bozkir A, Devrim B. Design and Evaluation of Hydrophobic Ion-Pairing Complexation of Lysozyme with Sodium Dodecyl Sulfate for Improved Encapsulation of Hydrophilic Peptides/Proteins by Lipid/Polymer Hybrid Nanoparticles. J Nanomed Nanotechnol 6, (2015).

15. Beach, M. A. et al. Polymeric Nanoparticles for Drug Delivery. Chem. Rev. 124, 5505–5616 (2024).

16. Meka, V. S. et al. A comprehensive review on polyelectrolyte complexes. Drug Discovery Today 22, 1697–1706 (2017).

17. Murthy, S. K. Nanoparticles in modern medicine: state of the art and future challenges. Int J Nanomedicine 2, 129–141 (2007).

18. Williams, J. A., Meynell, M. J. & Watson, A. B. Benethamine penicillin; a study of its use in a clinic for septic hands. Br Med J 1, 716–718 (1956).

19. Silva, E. L. et al. Nanostructured lipid carriers loaded with tributyrin as an alternative to improve anticancer activity of all-trans retinoic acid. Expert Rev Anticancer Ther 15, 247– 256 (2015).

20. Cui, S. et al. Correlation of the cytotoxic effects of cationic lipids with their headgroups. Toxicol Res (Camb) 7, 473–479 (2018).

21. Wibel, R., Friedl, J. D., Zaichik, S. & Bernkop-Schnürch, A. Hydrophobic ion pairing (HIP) of (poly)peptide drugs: Benefits and drawbacks of different preparation methods. European Journal of Pharmaceutics and Biopharmaceutics 151, 73–80 (2020).

22. Zheng, L., Bandara, S. R., Tan, Z. & Leal, C. Lipid nanoparticle topology regulates endosomal escape and delivery of RNA to the cytoplasm. Proceedings of the National Academy of Sciences 120, e2301067120 (2023).

23. Leung, A. K. K. et al. Lipid Nanoparticles Containing siRNA Synthesized by Microfluidic Mixing Exhibit an Electron-Dense Nanostructured Core. J. Phys. Chem. C 116, 18440– 18450 (2012).

24. Gigante, A. et al. Non-viral transfection vectors: are hybrid materials the way forward? Med. Chem. Commun. 10, 1692–1718 (2019).

25. Knapp, C. M., Guo, P. & Whitehead, K. A. Lipidoid Tail Structure Strongly Influences siRNA Delivery Activity. Cellular and Molecular Bioengineering 9, 305–314 (2016).

26. Ow, Y.-L. P., Green, D. R., Hao, Z. & Mak, T. W. Cytochrome c: functions beyond respiration. Nature Reviews Molecular Cell Biology 9, 532–542 (2008).

27. Kozuch, D. J., Ristroph, K., Prud’homme, R. K. & Debenedetti, P. G. Insights into Hydrophobic Ion Pairing from Molecular Simulation and Experiment. ACS Nano 14, 6097– 6106 (2020).

28. Sun, S. et al. pH-sensitive poly(lactide-co-glycolide) nanoparticle composite microcapsules for oral delivery of insulin. Int J Nanomedicine 10, 3489–3498 (2015).

29. Ristroph, K. et al. Internal liquid crystal structures in nanocarriers containing drug hydrophobic ion pairs dictate drug release. Journal of Colloid and Interface Science 582, 815–824 (2021).

30. Atef, B., Ishak, R. A. H., Badawy, S. S. & Osman, R. Exploring the potential of oleic acid in nanotechnology-mediated dermal drug delivery: An up-to-date review. Journal of Drug Delivery Science and Technology 67, 103032 (2022).

31. Yener, G., Gönüllü, U., Uner, M., Değim, T. & Araman, A. Effect of vehicles and penetration enhancers on the in vitro percutaneous absorption of celecoxib through human skin. Pharmazie 58, 330–333 (2003).

32. Salimi, A., Emam, M. & Mohammad Soleymani, S. Increase adapalene delivery using chemical and herbal enhancers. J Cosmet Dermatol 20, 3011–3017 (2021).

33. Murakami, T. et al. Topical delivery of keloid therapeutic drug, tranilast, by combined use of oleic acid and propylene glycol as a penetration enhancer: evaluation by skin microdialysis in rats. J Pharm Pharmacol 50, 49–54 (1998).

34. Griesser, J. et al. Hydrophobic ion pairing: Key to highly payloaded self-emulsifying peptide drug delivery systems. International Journal of Pharmaceutics 520, 267–274 (2017).

35. Gray, M. A. et al. Intracellular Biomacromolecule Delivery by Stimuli-Responsive Protein Vesicles Loaded by Hydrophobic Ion Pairing. ACS Omega (2025) doi:10.1021/acsomega.4c07666.

36. Da Silva Sanchez, A.J. et al. Universal Barcoding Predicts In Vivo ApoE-Independent Lipid Nanoparticle Delivery. Nano Lett. 22, 4822–4830 (2022).

37. Zhao, N. et al. Transfecting the hard-to-transfect lymphoma/leukemia cells using a simple cationic polymer nanocomplex. Journal of controlled release 159, 104–110 (2012).

38. Turtle, C. J. et al. CD19 CAR–T cells of defined CD4+: CD8+ composition in adult B cell ALL patients. The Journal of clinical investigation 126, 2123–2138 (2016).

39. Porter, D. L., Levine, B. L., Kalos, M., Bagg, A. & June, C. H. Chimeric antigen receptor– modified T cells in chronic lymphoid leukemia. New England Journal of Medicine 365, 725– 733 (2011).

40. Wang, Z., Wu, Z., Liu, Y. & Han, W. New development in CAR-T cell therapy. Journal of hematology & oncology 10, 1–11 (2017).

41. Maude, S. L. et al. Sustained Remissions with CD19-Specific Chimeric Antigen Receptor (CAR)-Modified T Cells in Children with Relapsed/Refractory ALL. (American Society of Clinical Oncology, 2016).

42. Yee Mon, K. J. et al. Functionalized nanowires for miRNA-mediated therapeutic programming of naïve T cells. Nature Nanotechnology 19, 1190–1202 (2024).

43. Mitchell, D. A. et al. Selective modification of antigen-specific T cells by RNA electroporation. Human gene therapy 19, 511–521 (2008).

44. Liu, L., Johnson, C., Fujimura, S., Teque, F. & Levy, J. A. Transfection optimization for primary human CD8+ cells. Journal of immunological methods 372, 22–29 (2011).

45. Karra, D. & Dahm, R. Transfection techniques for neuronal cells. J Neurosci 30, 6171–6177 (2010).

46. Ding, W. et al. Mechanism-Driven Technology Development for Solving the Intracellular Delivery Problem of Hard-To-Transfect Cells. Nano Lett. 23, 5859–5867 (2023).

47. Hoang, D. M. et al. Stem cell-based therapy for human diseases. Signal Transduction and Targeted Therapy 7, 272 (2022).

48. Pardi, N. et al. Expression kinetics of nucleoside-modified mRNA delivered in lipid nanoparticles to mice by various routes. Journal of Controlled Release 217, 345–351 (2015).

49. Melo, M. et al. Immunogenicity of RNA replicons encoding HIV Env immunogens designed for self-assembly into nanoparticles. Molecular Therapy 27, 2080–2090 (2019).

50. Melamed, J. R. et al. Ionizable lipid nanoparticles deliver mRNA to pancreatic β cells via macrophage-mediated gene transfer. Science Advances 9, eade1444.

51. Biscans, A. et al. Diverse lipid conjugates for functional extra-hepatic siRNA delivery in vivo. Nucleic Acids Research 47, 1082–1096 (2019).

52. Kang, J.-Y. et al. Engineered small extracellular vesicle-mediated NOX4 siRNA delivery for targeted therapy of cardiac hypertrophy. J Extracell Vesicles 12, e12371 (2023).

53. Sugo, T. et al. Development of antibody-siRNA conjugate targeted to cardiac and skeletal muscles. J Control Release 237, 1–13 (2016).

54. Slaughter, K. V. et al. Ionizable Drugs Enable Intracellular Delivery of Co-Formulated siRNA. Advanced Materials n/a, 2403701 (2024).

55. Papakyriakopoulou, P., Valsami, G. & Kadoglou, N. P. E. Nose-to-Heart Approach: Unveiling an Alternative Route of Acute Treatment. Biomedicines 12, (2024).

56. Harris, E., Zimmerman, D., Warga, E., Bamezai, A. & Elmer, J. Nonviral gene delivery to T cells with Lipofectamine LTX. Biotechnology and Bioengineering 118, 1674–1687 (2021).

57. Schaible, P., Bethge, W., Lengerke, C. & Haraszti, R. A. RNA Therapeutics for Improving CAR T-cell Safety and Efficacy. Cancer Res 83, 354–362 (2023).

58. Chen, L.-Y. et al. Successful application of anti-CD19 CAR-T therapy with IL-6 knocking down to patients with central nervous system B-cell acute lymphocytic leukemia. Translational oncology 13, 100838 (2020).

59. Teoh, P. J. & Chng, W. J. CAR T-cell therapy in multiple myeloma: more room for improvement. Blood Cancer Journal 11, 84 (2021).

60. Lee, Y.-H. et al. PD-1 and TIGIT downregulation distinctly affect the effector and early memory phenotypes of CD19-targeting CAR T cells. Molecular Therapy 30, 579–592 (2022).

61. Sultana, N. et al. Optimizing Cardiac Delivery of Modified mRNA. Molecular Therapy 25, 1306–1315 (2017).

62. Evers, M. J. et al. Delivery of modified mRNA to damaged myocardium by systemic administration of lipid nanoparticles. Journal of Controlled Release 343, 207–216 (2022).

63. Turnbull, I. C. et al. Myocardial delivery of lipidoid nanoparticle carrying modRNA induces rapid and transient expression. Molecular Therapy 24, 66–75 (2016).

64. Soroudi, S., Jaafari, M. R. & Arabi, L. Lipid nanoparticle (LNP) mediated mRNA delivery in cardiovascular diseases: Advances in genome editing and CAR T cell therapy. Journal of Controlled Release 372, 113–140 (2024).

65. Dautel, D. R. & Champion, J. A. Protein Vesicles Self-Assembled from Functional Globular Proteins with Different Charge and Size. Biomacromolecules 22, 116–125 (2021).

