## Supplemental Information for "Hydrophobic Ion Pairing for Simple, Non-Toxic Transfection"

**MATERIALS**

Benethamine (*N*-benzyl-2-phenylthanamine) (catalog number: APOH316Cde11), OA (catalog number: O07501), SD (catalog number: D4422), CytC (catalog number: 63103), fetal bovine serum (FBS), trypsin-EDTA, and imidazole were obtained from Sigma-Aldrich. Butanol was obtained from Millipore Sigma. Liptofectamine RNAiMAX Transfection Reagent (catalog number: 13778030) and siRNA silencing eGFP (catalog number: AM4626) were from Fisher Scientific. NIH 3T3 fibroblasts (GFP3T3) reporter cells (catalog number: AKR-214) were from CellBio Labs. Dulbecco’s modified Eagle’s medium (DMEM), human and mouse anti-LAMP1 siRNA (s80802), and anti-LAMP1 antibody (53-1079-42) were purchased from ThermoFisher. Luciferase encoding pDNA was purchased from Aldevron (5001).

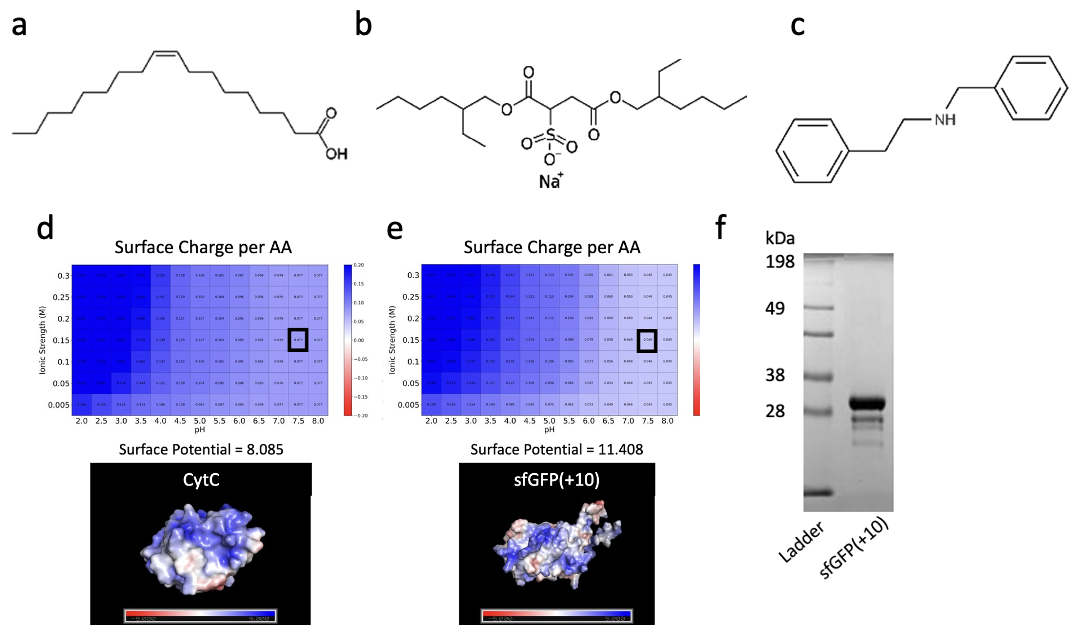

**Figure S1.** Characterization of counterions and proteins. Counterion structures of (a) OA with a LogP = 6.78, (b) SD with a LogP = 5.2, and (c) benethamine with a LogP = 3.6^1^. Surface potential of (d) Cytochrome C and (e) sfGFP(+10) per amino acid determined by using the protein sequence to predict structure^2^, electrostatics calculations^3^, and surface analysis^4^. (f) SDS-PAGE gels of sfGFP(+10).

| **Cargo : Counterion** | **Mass Ratio** | **Molar Ratio** | **Charge Ratio** |
| --- | --- | --- | --- |
| sfGFP(+10) : OA | 1 : 4 | 1 : 396 | 1 : 35 |
| sfGFP(+10) : SD | 1 : 1 | 1 : 62 | 1 : 5.4 |
| CytC : OA | 1 : 3 | 1 : 127 | 1 : 16 |
| CytC : SD | 1 : 4 | 1 : 108 | 1 : 13 |
| siRNA : BA | 1 : 4 | 1 : 0.585 | 1 : 0.03 |
| pDNA : BA | 1 : 15 | 1 : 0.008 | 1 : 1.6 $\times{10}^{-6}$ |

**Table S1**. HIP protein complex mass, molar, and charge ratios using oleic acid (OA), sodium docusate (SD), and benethamine (BA). Charge ratios were calculated based on charged surface accessible amino acid residues in cargo for proteins or based on number of base pairs for nucleic acids.

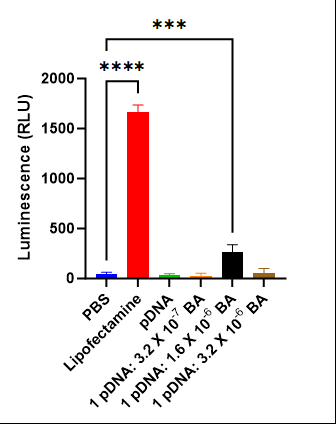

**Figure S2**. Plasmid DNA (pDNA) delivery using HIP. Quantification of luciferase encoded pDNA HIP delivery using a luciferase assay in HeLa cells 72 hours after treatment. HIP groups were composed of equal volumes of 1 mg/ml pDNA with 15 mg/ml of benethamine (BA) and were incubated immediately after HIP mixing. pDNA was mixed with an equal volume of 1X PBS because counterions were dissolved in 1X PBS. 2µg pDNA was dosed in the presence of FBS for all groups except for Lipofectamine™ 2000 group. pDNA was mixed with counterion, then mixed with FBS containing media and administered to cells. *** p < 0.001, and **** p < 0.0001.

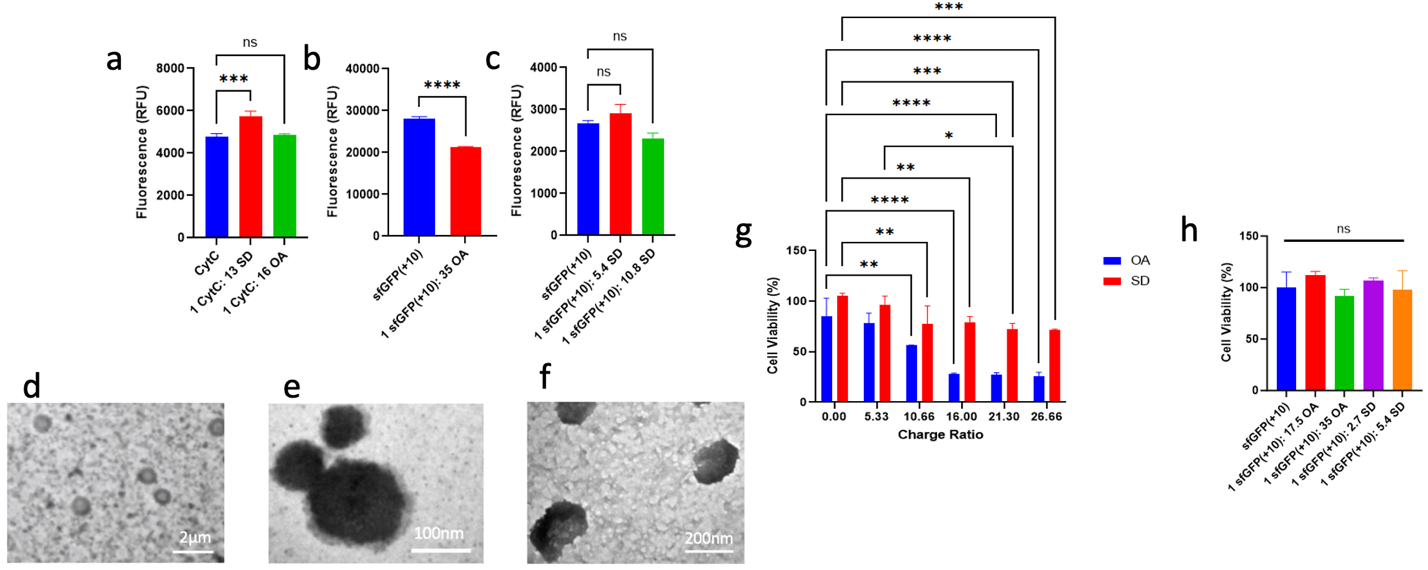

**Figure S3.** HIP protein delivery. Counterion influence on cargo fluorescence as an indicator of change in protein structure for a) CytC, b) sfGFP(+10) with OA and c) sfGFP(+10) with SD. TEM images of HIP complexes observed at charge ratios of d) 1 CytC: 1.8 OA, e) 1 CytC: 5.3 OA, and f) 1 CytC: 16 OA. Influence on HeLa cell viability after 48 hours incubation in 10% FBS supplemented media with of g) 20 µM cytotoxic Cytochrome C or h) non-toxic sfGFP(+10) in HIP complexes with OA or SD at given charge ratios measured by MTT assay. Significant decrease in viability begins at a charge ratio of 10.66 for both counterions, indicating the minimum required charge ratio for successful cytosolic CytC delivery. Non-toxic sfGFP(+10) cargo does not exhibit reduced viability, indicating it is CytC and not the hydrophobic ions responsible for the cytotoxic effect seen in g). * p < 0.05, ** p < 0.01, *** p < 0.001, and **** p < 0.0001.

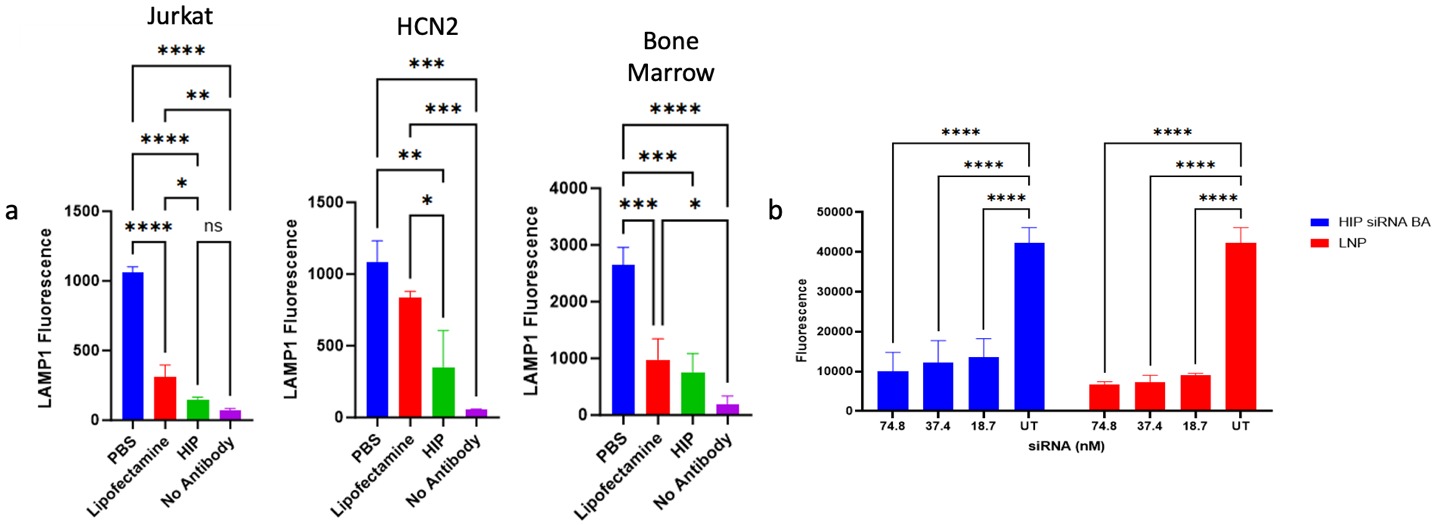

**Figure S4.** Fluorescence data of HIP siRNA delivery of a) hard-to-transfect cell types (Jurkat, HCN2, and bone marrow) and b) LNP comparison delivery in NIH3T3/eGFP cells. 75 nM siRNA, or varying amounts of siRNA as noted was given in the presence of FBS for all groups except for Lipofectamine™ groups and treated for 48 hours. Gene silencing a) was measured by labelling cells with either a human or mouse anti-LAMP1 antibody and measuring mean fluorescence using flow cytometry and b) using plate reader GFP fluorescence.

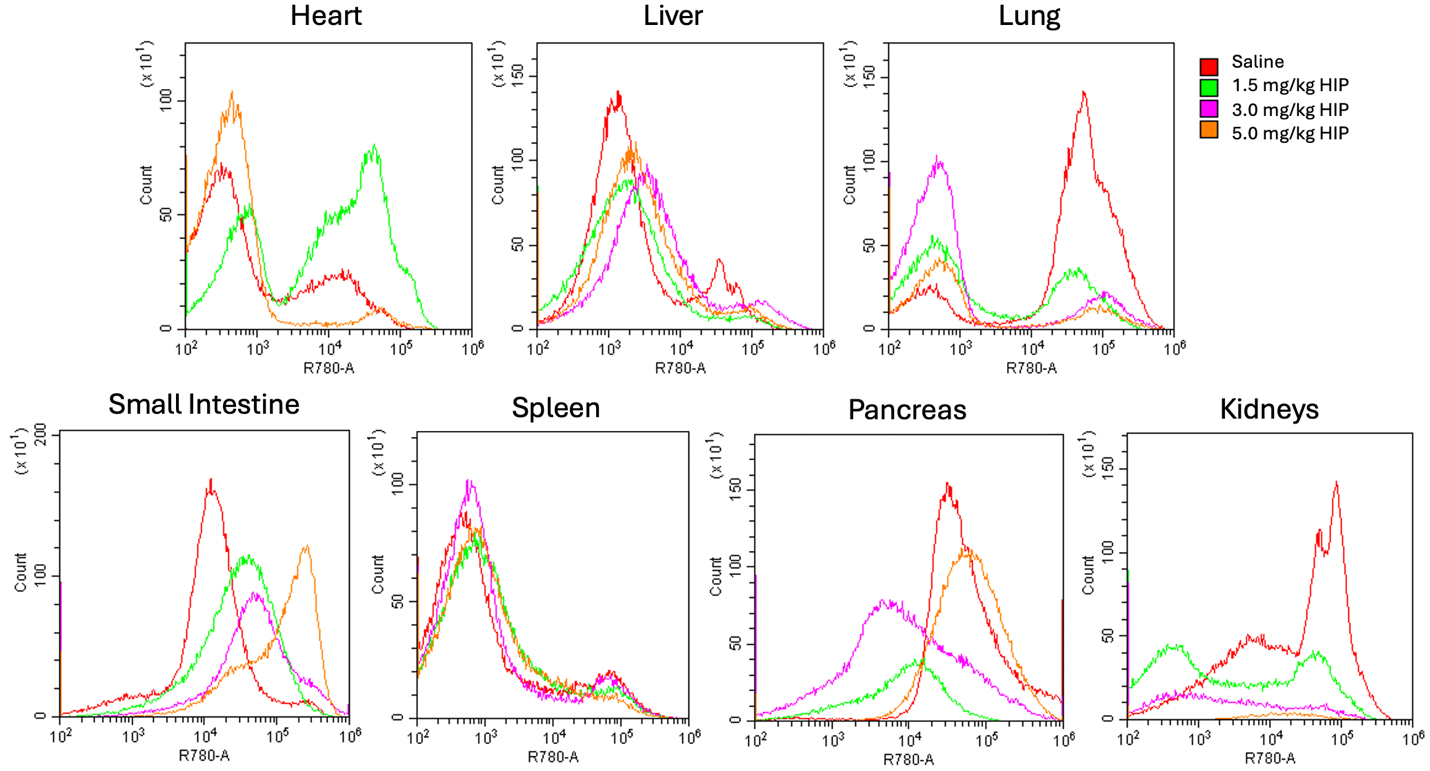

**Figure S5**. Preliminary IP *in vivo* biodistribution to select anti-LAMP1 siRNA dose (1.5, 3.0, or 5.0 mg/kg) and organs of interest for HIP intracellular delivery after 48 hours in BALB/c (n=2) mice. Organs were homogenized, red blood cells were removed, and an anti-LAMP1 antibody was used to assess LAMP1 gene silencing using flow cytometry. Data from single mouse shown as representative.

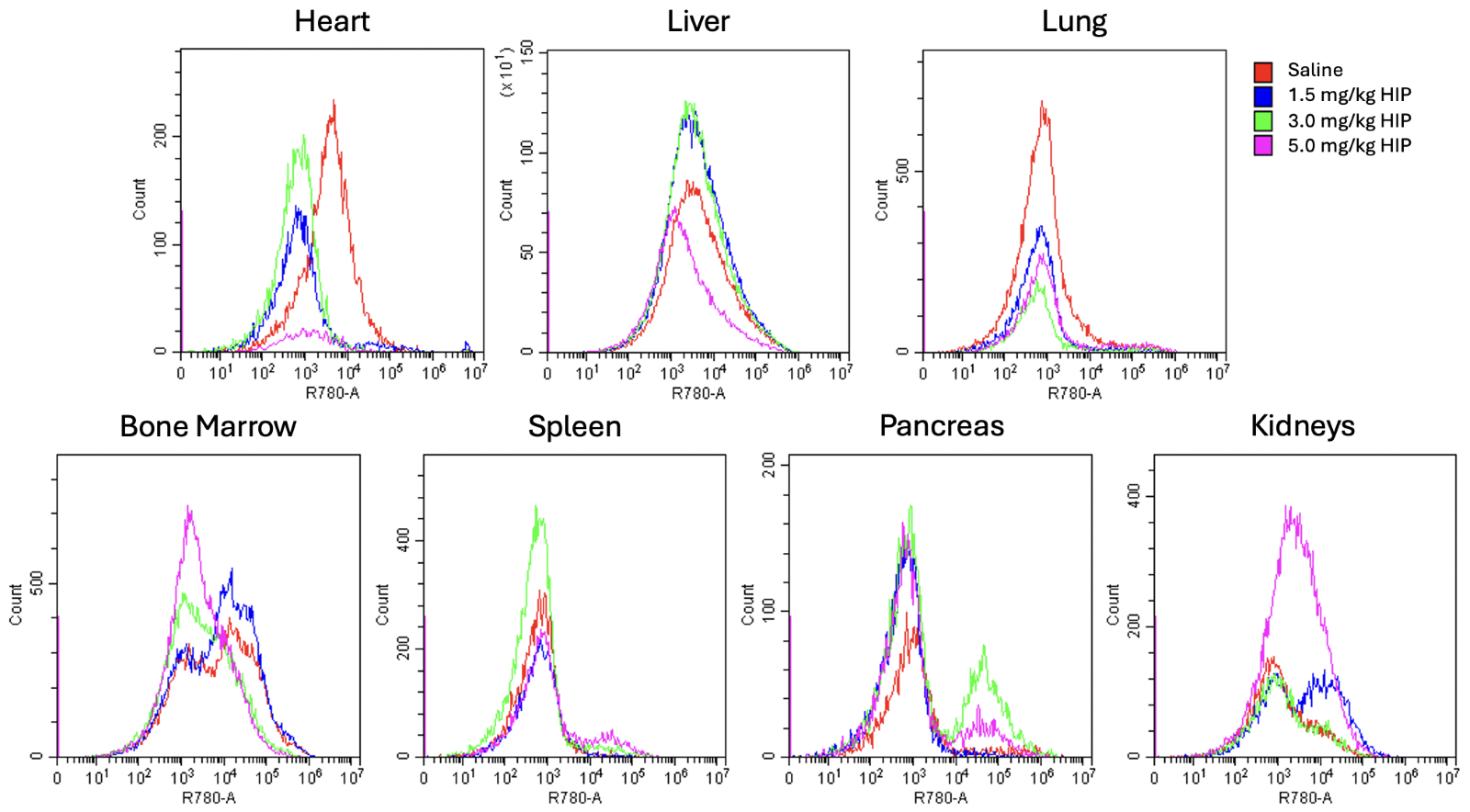

**Figure S6.** Preliminary IV *in vivo* biodistribution to select anti-LAMP1 siRNA dose (1.5, 3.0, or 5.0 mg/kg) and organs of interest for HIP intracellular delivery after 48 hours in BALB/c (n=2) mice. Organs were homogenized, red blood cells were removed, and an anti-LAMP1 antibody was used to assess LAMP1 gene silencing using flow cytometry. Data from single mouse shown as representative.

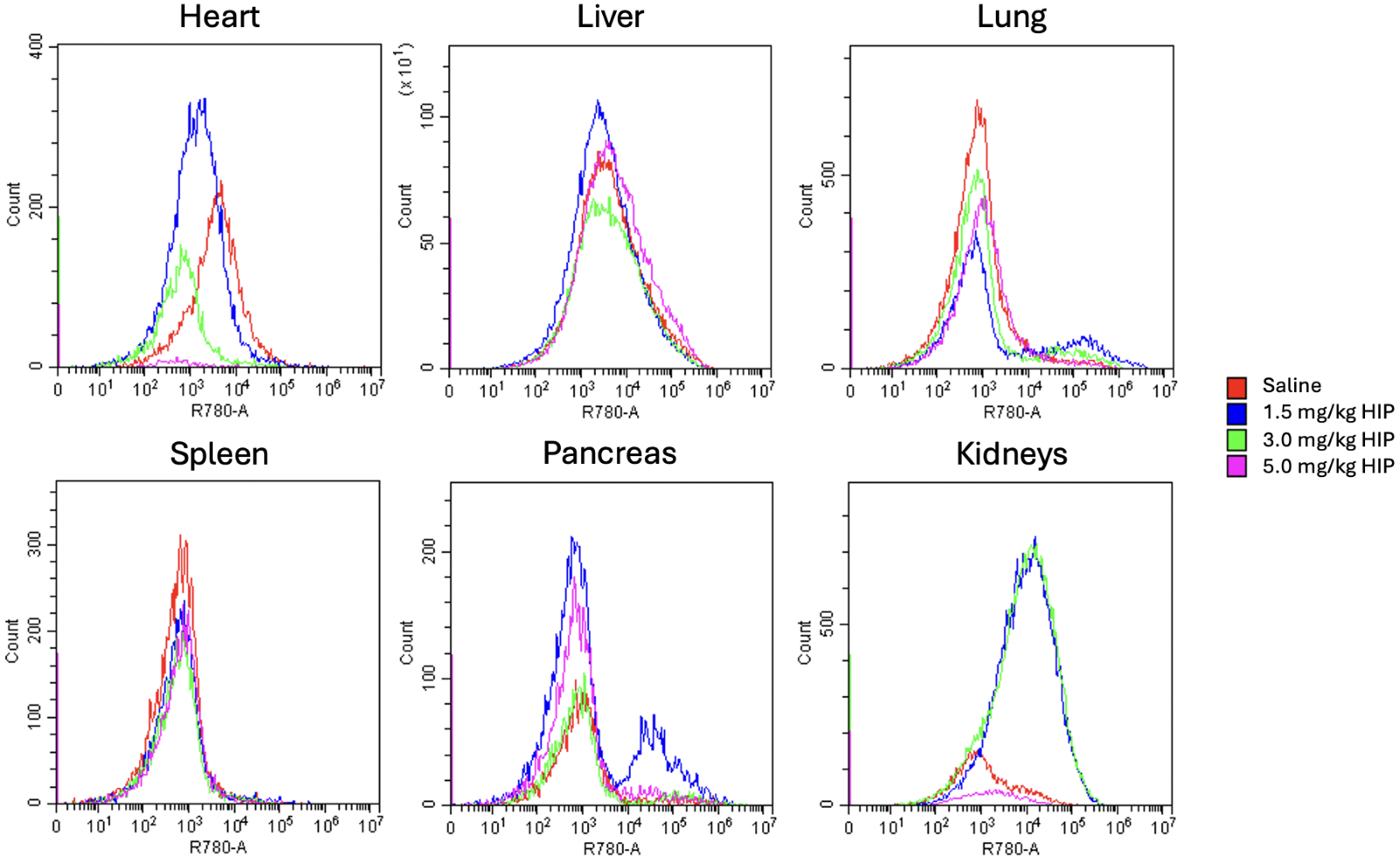
**Figure S7.** Preliminary IN *in vivo* biodistribution to select anti-LAMP1 siRNA dose (1.5, 3.0, or 5.0 mg/kg) and organs of interest for HIP intracellular delivery after 48 hours in BALB/c (n=2) mice. Organs were homogenized, red blood cells were removed, and an anti-LAMP1 antibody was used to assess LAMP1 gene silencing using flow cytometry. Data from single mouse shown as representative.

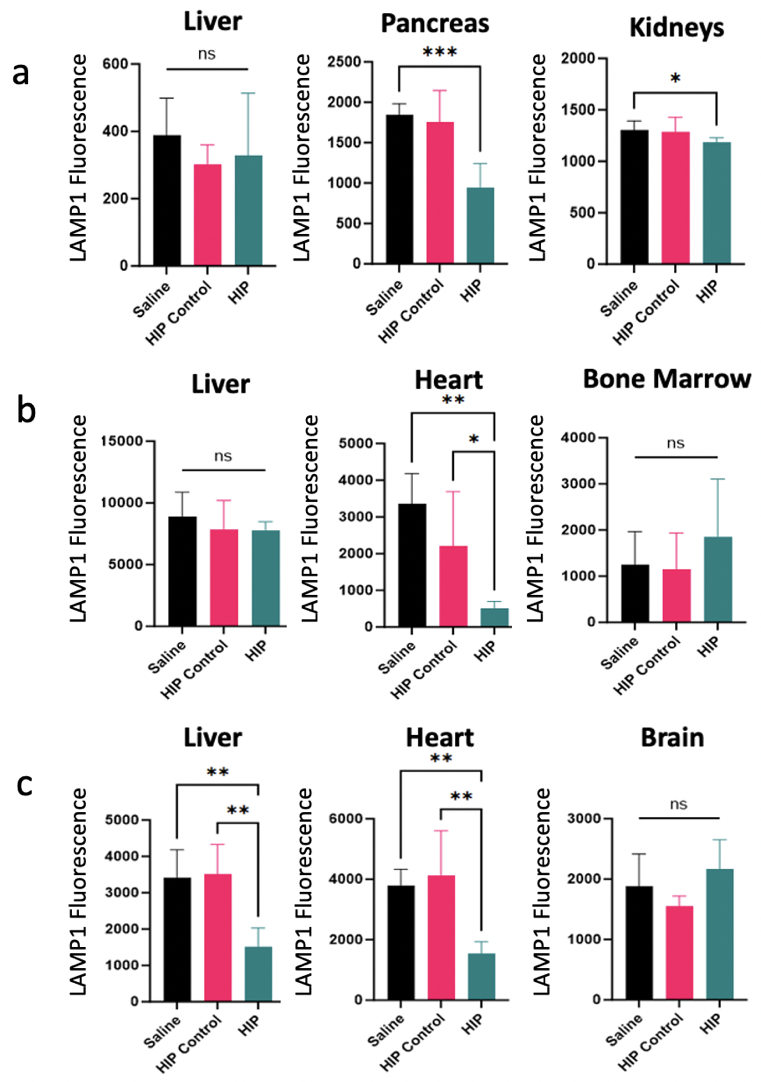

**Figure S8.** *In vivo* functional delivery biodistributions using a) intraperitoneal (IP), b) intravenous (IV), and c) intranasal (IN) routes in (n=5) BALB/c mice using an anti-LAMP1 siRNA dose of 1.5 mg/kg, 5 mg/kg, and 1.5 mg/kg, respectively. Organs were homogenized, red blood cells were removed, and an anti-LAMP1 antibody was used to assess LAMP1 surface expression using flow cytometry 48 hours after treatment. One-way ANOVA and the F-test were used to compare groups for statistical significance. Outliers were removed (from saline group only as experiments were performed on different days) using Grubbs’ statistical test. * p < 0.05, ** p < 0.01, and *** p < 0.001.

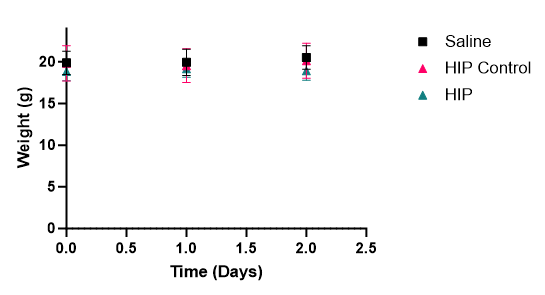

**Figure S9.** Mouse weight following HIP-siRNA complex administration at day 0.

**References**

(1)	Ristroph, K. D.; Prud’homme, R. K. Hydrophobic Ion Pairing: Encapsulating Small Molecules, Peptides, and Proteins into Nanocarriers. *Nanoscale Adv.* **2019**, *1* (11), 4207–4237. https://doi.org/10.1039/C9NA00308H.

(2)	Mirdita, M.; Schütze, K.; Moriwaki, Y.; Heo, L.; Ovchinnikov, S.; Steinegger, M. ColabFold: Making Protein Folding Accessible to All. *Nature Methods* **2022**, *19* (6), 679–682. https://doi.org/10.1038/s41592-022-01488-1.

(3)	Dolinsky, T. J.; Nielsen, J. E.; McCammon, J. A.; Baker, N. A. PDB2PQR: An Automated Pipeline for the Setup of Poisson-Boltzmann Electrostatics Calculations. *Nucleic Acids Res* **2004**, *32* (Web Server issue), W665-667. https://doi.org/10.1093/nar/gkh381.

(4)	Hebditch, M.; Warwicker, J. Web-Based Display of Protein Surface and pH-Dependent Properties for Assessing the Developability of Biotherapeutics. *Scientific Reports* **2019**, *9* (1), 1969. https://doi.org/10.1038/s41598-018-36950-8.
